# An Interpretable Performance Metric for Auditory Attention Decoding Algorithms in a Context of Neuro-Steered Gain Control

**DOI:** 10.1101/745695

**Authors:** Simon Geirnaert, Tom Francart, Alexander Bertrand

## Abstract

In a multi-speaker scenario, a hearing aid lacks information on which speaker the user intends to attend, and therefore it often mistakenly treats the latter as noise while enhancing an interfering speaker. Recently, it has been shown that it is possible to decode the attended speaker from the brain activity, e.g., recorded by electroencephalography sensors. While numerous of these auditory attention decoding (AAD) algorithms appeared in the literature, their performance is generally evaluated in a non-uniform manner. Furthermore, AAD algorithms typically introduce a trade-off between the AAD accuracy and the time needed to make an AAD decision, which hampers an objective benchmarking as it remains unclear which point in each algorithm’s trade-off space is the optimal one in a context of neuro-steered gain control. To this end, we present an interpretable performance metric to evaluate AAD algorithms, based on an adaptive gain control system, steered by AAD decisions. Such a system can be modeled as a Markov chain, from which the minimal expected switch duration (MESD) can be calculated and interpreted as the expected time required to switch the operation of the hearing aid after an attention switch of the user, thereby resolving the trade-off between AAD accuracy and decision time. Furthermore, we show that the MESD calculation provides an automatic and theoretically founded procedure to optimize the number of gain levels and decision time in an AAD-based adaptive gain control system.

## I. Introduction

Current hearing aids and cochlear implants have major difficulties in reducing background noise in a so-called ‘cocktail party’ scenario, in which multiple speakers talk simultaneously. It is however known that the human brain is capable of ‘filtering’ out the attended speaker and ignoring all competing speakers [1]. State-of-the-art noise reduction algorithms are very well able to extract a single speech source and subtract background noise or interfering speakers as well, but a fundamental problem is that the hearing aid should decide which speaker is the attended speaker (i.e., the speaker a user intends to attend) and which other speakers should be treated as noise sources. Currently, this is done using unreliable heuristics based on, e.g., speaker intensity or look direction.

Recently, it has been demonstrated that the attended speech signal can be decoded from cortical brain responses and that its dynamical changes are tracked by the brain [2]. More specifically, it is shown that the brain tracks the envelope of the attended speech signal [3]–[5]. These advances have led to a multitude of algorithms that decode attention from the brain using magneto- or electroencephalography (EEG) (e.g., [6]–[11]). These auditory attention decoding (AAD) algorithms are paramount to design ‘neuro-steered’ hearing aids. To this end, a first full modular pipeline was presented in [12], where AAD was used in combination with source separation and noise reduction algorithms on a hearing aid’s microphone recordings. Later, also alternative pipelines were presented based on other source separation algorithms [13]–[16]. Furthermore, the effects of different boundary conditions, such as speaker positions or noisy and reverberant conditions, have already extensively studied as well (e.g., [17]–[19]).

However, an important question is how these AAD algorithms should be evaluated. Their accuracy, measured as the percentage of decision windows in which the attention was decoded correctly, depends on the length of the decision window, which defines how much EEG data are available to make a decision. Because of the low signal-to-noise ratio (SNR) of the neural response to the speech signals in the EEG, the accuracy increases with the length of the decision window. However, a longer decision window implies that the algorithm also needs more time to, e.g., react on a switch in attention, which results in a trade-off. This trade-off between accuracy and decision window length results in three fundamental issues regarding the evaluation of AAD algorithms:

- The dependence of the accuracy on the decision window length hinders easy statistical comparison, as the different decision window lengths need to be taken into account as an extra factor. This hampers drawing adequate statistical conclusions.
- Algorithm A might perform better than algorithm B for smaller decision window lengths, while algorithm B might perform better than algorithm A for large decision window lengths, leading to inconclusiveness when benchmarking both algorithms.
- In several scientific reports, only one decision window length with corresponding accuracy is reported. A different choice of the decision window lengths (e.g., across two scientific reports) then obstructs a fair comparison.

The aforementioned problems motivate the need for a single-number metric to capture the overall AAD performance, which also takes the trade-off between accuracy and decision time into account by selecting the optimal point on the trade-off curve that is the most relevant in the context of adaptive gain control for neuro-steered hearing aids.

In [7], the Wolpaw information transfer rate (ITR_W_) 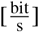 is adopted from the brain-computer interfacing (BCI) community to combine the accuracy and the decision window length in a single metric as follows [20]:

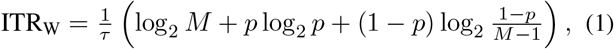

with *p* the accuracy (probability of a correct decision), *τ* the decision time (here: decision window length) and *M* the number of classes (here: speakers). Similarly, in [21], the Nykopp ITR (ITR_N_) is used to evaluate AAD algorithms, which assumes an adaptive brain-computer interface setting in which not every time a decision has to be made [21]. The ITR_W_ was originally defined to quantify the performance of BCI systems that are used to re-establish or enhance communication and control for paralyzed individuals with severe motor impairments [20]. It quantifies the number of bits that can be transferred per time unit and matches as such the specific context of communicating through brain waves. However, the ITR_W/N_ has no such clear interpretation in the context of AAD for neuro-steered hearing prostheses and is therefore not per se a relevant criterion to compare AAD algorithms. Instead, we are interested in *how fast* a hearing aid can switch its operation from one speaker to another, following an intentional attention switch of the user, based on consecutive AAD decisions and taking into account that some decisions may be incorrect.

The lack of a *interpretable* metric in the context of neurosteered hearing prostheses, which combines both decision time and accuracy in a single metric, and which facilitates making unambiguous conclusions on performance and easy comparisons between algorithms, motivates the design of a new metric, which we refer to as the *minimal expected switch duration* (MESD)^1^. The MESD metric is based on the performance of an adaptive gain control system that is optimized for the AAD algorithm under test. Therefore, the derivation of the MESD metric also leads to an automatic and theoretically founded procedure to optimize the step size and decision frequency in an AAD-based adaptive gain control system, thereby avoiding a tedious manual tuning.

In Section II, we develop this new metric step-by-step, leading to a closed-form expression based on which the metric can be computed. In Section III, we give examples of the MESD metric on real EEG/audio data, as well as a comparison with the ITR_W/N_ metric. Conclusions are drawn in Section IV.

### Disclaimer

A conference precursor of this manuscript has been published in [22]. The current manuscript provides a more accessible introduction to the MESD metric, provides the complete mathematics (including all the derivations and proofs that were missing in [22], as well as a formal algorithm description), contains a more extensive and more quantitative validation (including statistical analyses), contains an extensive comparison with other performance metrics, and includes more details on the hyperparameter choice.

## II. Expected switch duration

### A. An adaptive gain control system

Given that AAD algorithms decode the attention of a hearing aid user, hearing aids could benefit from an adaptive gain/volume control system. Given a two-speaker situation, such a system would allow to adaptively over time change the gain of speaker one versus speaker two, tracking the attention of the hearing aid user (see Fig. 1). We, however, want to avoid the usage of only two volume settings or gain control ‘states’, i.e., all-or-nothing amplification of both speakers, as this would cause perceptually unpleasant spurious and sudden switching of speakers (of which many by mistake). Moreover, we want to enable the user to adequately react when the system starts switching towards the wrong speaker due to AAD errors, before the attended speaker becomes unintelligible. As a result, the system should have many states to gradually and adaptively change the relative gain between both speakers.

**Fig. 1:**
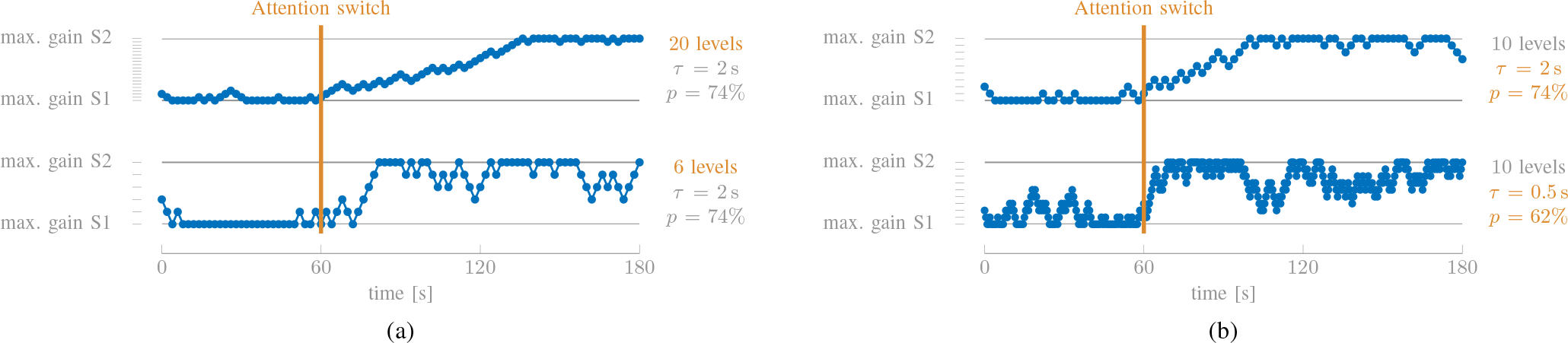
This example illustrates the two fundamental issues regarding an adaptive gain control system with decision window length *τ* and AAD accuracy *p*. In the first minute, speaker one (S1) is the attended speaker, while after 60 s, the attention switches to speaker two (S2). (a) When the number of gain levels decreases, the gain switch is performed faster, but the overall gain process is less stable. (b) Decreasing the decision window length - and correspondingly the accuracy - results in a faster gain switch, but less stable gain process, and vice versa.

However, this results in two crucial design parameters which both affect the performance of the system, each leading to a fundamental trade-off, which is illustrated in Fig. 1:

1. *How many gain levels should we use?* As Fig. 1a illustrates, using fewer gain levels results in a faster gain switch after an attention switch, but also results in a less stable gain process, negatively affecting the comfort of the user. Increasing the number of gains stabilizes the gain process and thus results in a more robust gain control, but increases the gain switch time.
2. *How often should we take a step?* A short decision window length corresponds to a fast gain control system - as less EEG and audio data need to be buffered before a decision can be made - and thus a fast gain switch (Fig. 1b). However, as is indicated in Section I, a shorter decision window length also corresponds to a lower accuracy, resulting in a more unstable gain process - vice versa for a longer decision window length.

Note that optimizing a *discrete* gain level system does not imply that there needs to be a discrete implementation in a hearing aid. One could as well continuously interpolate between the discrete gain levels to provide a more pleasant user experience. In that case, optimizing the rate of change of the volume (e.g., the slope) corresponds to optimizing the number of gain levels.

In the following sections, we translate this adaptive gain control system into mathematics using a Markov chain model. This mathematical formulation will allow us to rigorously address these fundamental issues and optimize these two design parameters, as well as provide a way to properly evaluate and rank different AAD algorithms through the novel MESD metric, which is derived from the optimal gain control design for the AAD algorithm under test. This MESD metric is formally defined in Section II-E.

### B. Markov chain model

The adaptive gain control system of Section II-A can be straightforwardly translated into a mathematical model using a Markov chain (Fig. 2). Table I shows how the parameters of the Markov chain embody several concepts of the adaptive gain control system.

**Fig. 2:**
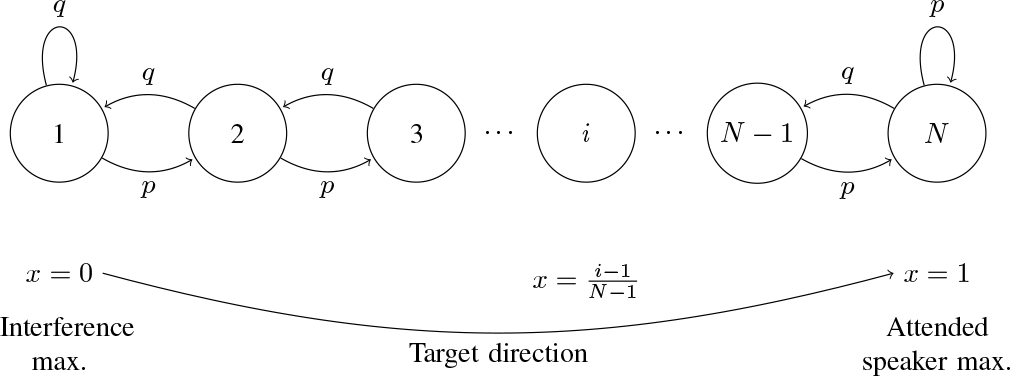
An adaptive gain control system can be modeled as a Markov chain with *N* states (gains) and a transition probability *p* in the target direction (attended speaker) equal to the accuracy of the AAD algorithm.

**TABLE I:** The different concepts of an adaptive gain control system have a straightforward translation to a Markov chain parameter.

| Adaptive gain control parameter | Markov chain parameter |
| --- | --- |
| gains | states $x \in [0, 1]$ |
| number of (relative) gain levels | number of states $N$ |
| AAD accuracy | transition probability $p$ |
| decision window length | step time $\tau$ |

The Markov chain contains *N* states, each corresponding to a relative gain level *x* ∈ [0, 1] of the attended speaker versus background noise, including the interfering speaker(s). For illustrative purposes throughout the manuscript, but without loss of generality, we will consider the example of a noiseless two-speaker scenario. In this case, *x* = 1 would correspond to a target relative amplification of the attended speaker versus the unattended speaker, which is typically constrained to still enabling the listener to switch attention to the other speaker. *x* = 0 then corresponds to the maximal suppression of the attended speaker with a similar constraint, while *x* = 0.5 implies equal gain for both speakers. These gain levels are assumed to be uniformly distributed over [0, 1], resulting in a one-to-one relation between state *i* and gain level *x*:

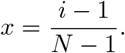

Given that *x* = 1 corresponds to the target gain level of the attended speaker, the transition probability *p* ∈ [0, 1] in the target direction is equal to the probability of a correct AAD decision, i.e., the AAD accuracy. Similarly, *q* = 1 − *p* corresponds to the probability of a wrong decision. In what follows, we assume that *p* > 0.5, i.e., the evaluated AAD algorithm performs at least better than chance level. A correct (step towards *x* = 1) or incorrect (step towards *x* = 0) decision always results in a transition to a neighboring state, except in state 1 or state *N*, where no state transition is made after an incorrect or correct decision, respectively (e.g., in state *N*, the gain is maximal for the attended speaker, which is the best the system can obtain). The latter is indicated by the self-loops in Fig. 2, which models the gain clipping in Fig. 1.

Each step takes *τ* seconds - the decision window length - as *τ* seconds of EEG and audio data need to be buffered before a new decision can be made. The application of an AAD algorithm on consecutive windows of *τ* seconds, which results in a gain process such as shown in Fig. 1, thus corresponds to a random walk process through the Markov chain. Note that the AAD accuracy *p* directly depends on this decision window length *τ*, as denoted before. The *p*(*τ*)-performance curve relates this AAD accuracy *p* with the decision window length *τ* for a particular AAD algorithm (see Fig. 4 for an example).

The two fundamental issues regarding the gain control system as listed in Section II-A, can now be translated into the optimization of the Markov chain parameters:

1. Optimizing the number of gain levels corresponds to the optimization of the number of states *N* (this will be derived in Section II-C).
2. Determining the time resolution with which the gain should be adapted corresponds to determining the step time *τ* (this will be derived in Section II-D). Note that equivalently, the transition probability *p* can be optimized. Addressing this second issue corresponds to jointly optimizing the AAD accuracy *p* and decision window length *τ*, as they are directly related through the *p*(*τ*)-performance curve. The resulting pair (*τ*_*opt*_, *p*_*opt*_) is called the *optimal working point* on the *p*(*τ*)-performance curve.

We will answer both of these questions through a mathematical analysis on the corresponding Markov chain in Section II-C and II-D, respectively, which will lead to the MESD metric in Section II-E. However, it should be emphasized that this Markov chain is a simplified model of a real gain control system and, as always, this mathematical tractability comes at the cost of making some simplifying assumptions. Indeed, a Markov chain assumes independence of the consecutive decisions^2^, which may be violated in a practical AAD algorithm, in particular when there is overlap in the data of consecutive windows.

### C. Optimizing the number of states N

We first optimize the number of states *N*, where we mainly target a stable gain process, tackling one of the trade-offs in Fig. 1 (left) (a stable gain process versus fast switching).

#### 1) Steady-state distribution

The steady-state distribution of the Markov chain in Fig. 2 is needed in order to analyze the behavior of the modeled adaptive gain control system. This steady-state distribution 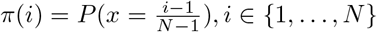 is defined as the probability to be in state *i* after an infinite number of random steps (starting from any position), for a fixed transition probability *p*. Defining 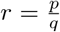, the steady-state distribution is shown in Appendix A to be equal to:

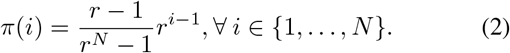

#### 2) P_0_-confidence interval

Based on the Markov chain model and the steady-state distribution, we determine a desirable operating region of the neuro-steered hearing aid via the *P*_0_-confidence interval 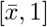. This is the smallest interval in which the system must operate for at least *P*_0_ percent of the time, despite the presence of AAD errors, while being in a steady-state regime. For example, if *P*_0_ = 0.8, we expect the hearing prosthesis to operate in the operating region 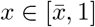 for at least 80% of the time. This implies that we search for the largest 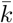 for which:

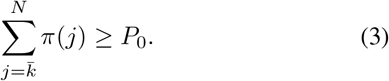

This leads to the following lower bound 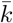 of the *P*_0_-confidence interval (the derivation is given in Appendix B):

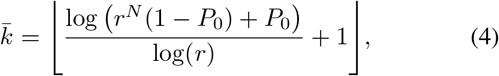

with [·] the flooring operation yielding an integer output. The resulting *P*_0_-confidence interval is thus defined as^3^:

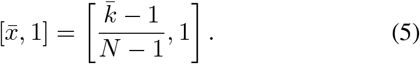

The *P*_0_-confidence interval is indicated in orange in Fig. 3.

**Fig. 3:**
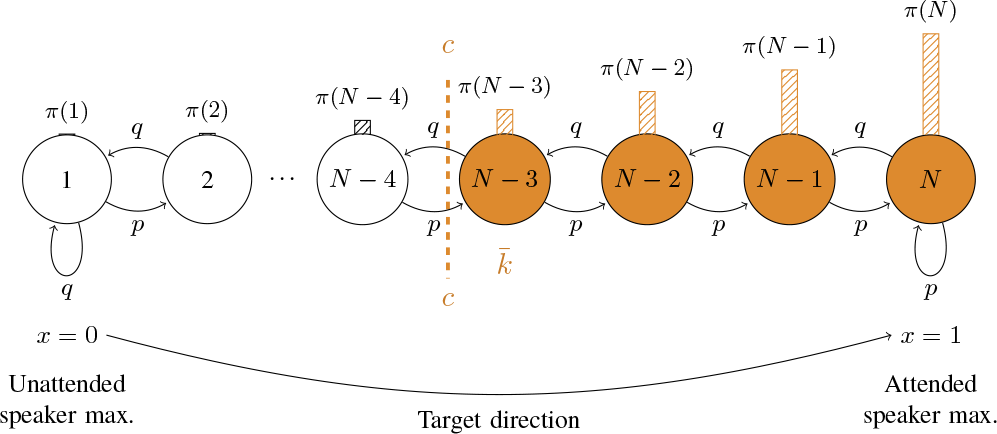
The *P*_0_-confidence interval in orange is the smallest set of states for which the sum of the steady-state probabilities (bars) is larger than *P*_0_. The second design constraint forces the lower bound 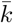 of this *P*_0_-confidence interval to be above a predefined level *c*, assuring stability of the system.

**Fig. 4:**
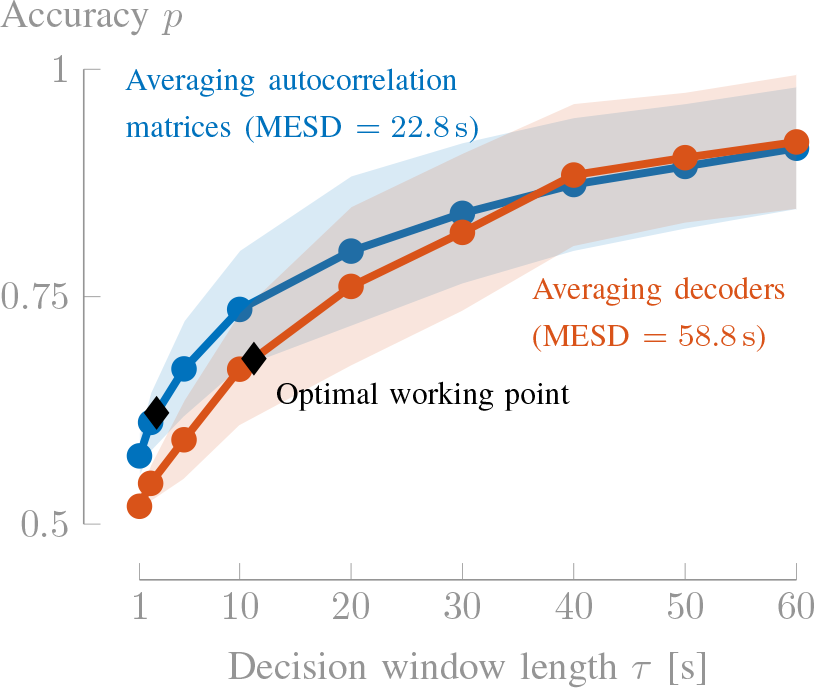
The MESD focuses on small decision window lengths as the relevant part of the performance curve, based on which it can be concluded that averaging autocorrelation matrices outperforms averaging of decoders.

#### 3) Design constraints

From Fig. 1, it can be intuitively seen that to minimize the gain switch duration, we have to minimize the number of states *N*. However, we also know that this conflicts with the stability of the gain process (Fig. 1). To guarantee a certain amount of stability or confidence of the system and comfort to the user, we propose the following design criteria for the Markov chain regarding *N*:

- The lower bound of the *P*_0_-confidence interval 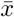 should be larger than a pre-defined ‘comfort level’ *c* that defines the target operating region, i.e., 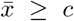. This comfort level *c* can be determined from hearing tests, for example, by interpreting it physiologically as the gain level below which it becomes uncomfortable to listen to the attended speaker (see Section III-A, where we will motivate to choose *c* = 0.65). By controlling *N*, we can thus ensure that the hearing prosthesis is in *P*_0_ percent of the time above this comfort level *c*, ergo, stabilizing the gain process. With (4) and (5), the above requirement results in the following inequality:

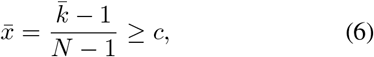

which should be viewed as a constraint when minimizing *N* (note that 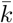 also depends on *N*). A key message here is that a *lower* accuracy *p* requires *more* states *N* in order to guarantee (6).
- *N* ≥ *N*_min_: a minimal number of states is desired to obtain a sufficiently smooth transition in the gain adaptation. In particular, we want to avoid the immediate crossing of the mid-level *x* = 0.5 (i.e., an immediate change of the loudest speaker) when leaving the *P*_0_-confidence interval due to an incorrect AAD decision. In cases where (6) is satisfied for *N* = 4, the *P*_0_-confidence interval also often^4^ contains state 3, which would result in an immediate crossing of *x* = 0.5 when leaving the *P*_0_-confidence interval due to an AAD error. Therefore, we propose to fix *N*_min_ = 5.

In practice, the minimal number of states *N* can be found by going over the candidate values *N* = *N*_min_ + *i*, with *i* = 0, 1, 2,…, in this specific order (as the gain switch duration increases with *N*), until a value *N* is found that satisfies (6). As shown in Appendix C, such a value of *N* can always be found, for any value of *c* and *P*_0_, assuming that *p* > 0.5.

### D. Finding the optimal working point (*τ*_*opt*_, *p*_*opt*_)

In Section II-C, we have constrained *N* such that the gain process has a minimum of stability, such that we can now focus on minimizing the gain switch time. In this section, we rigorously define the expected switch duration (ESD), which quantifies this gain switch time, and use it as a criterion to determine the optimal working point (*τ*_*opt*_, *p*_*opt*_).

#### 1) Mean hitting time

A fundamental metric within the Markov chain is the *mean hitting time* (MHT), which quantifies the expected number of steps *s* needed to arrive in target state *j* when starting from a given initial state *i*. The MHT is defined as:

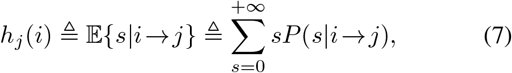

with *i*, *j* ∈ {1, …, *N*}, 𝔼 {·} denoting the expectation operator and where *P*(*s*∣*i* → *j*) is the probability that target state *j* is reached for the first time after *s* random steps, when starting in state *i*. Note that we are only interested in the MHT for the case where *i* ≤ *j*, i.e., when going from left to right in the Markov chain (Fig. 2). This corresponds to the case where the hearing aid switches from one speaker to the other. In Appendix D, we show that the MHT can be computed as:

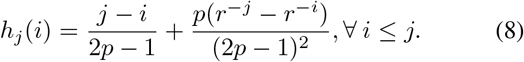

#### 2) Expected switch duration

We define a gain switch as the transition to the comfort level *c*, starting from *any* initial state *i* with a corresponding gain level outside the predefined working region [*c*, 1]. Note that this specific definition of a gain switch implies that we are aiming at quantifying the duration of a *stable* switch. The perceived gain switch towards the attended speaker by the hearing aid user would typically occur earlier, e.g., when *x* = 0.5 is reached. The corresponding gain switch time is called the *expected switch duration* (ESD) [s]. The ESD thus quantifies the time needed to change the operation of the system when the user shifts its attention *and* when the system is not yet in the desired operating region.

Assuming *k*_*c*_ is the first state corresponding to a relative gain *x* ≥ *c*:

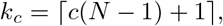

the ESD is formally defined as the expected time (step time *τ* times expected number of steps *s*) necessary to go from any state *i* < *k*_*c*_ to target state *k*_*c*_:

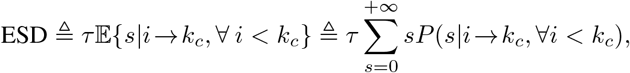

with *P*(*s*|*i* → *k*_*c*_, ∀ *i* < *k*_*c*_) the probability that target state *k*_*c*_ is reached for the first time after *s* steps, when starting from any state *i* < *k*_*c*_. Using marginalization in the initial state *i*, this can be written as:

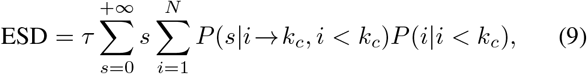

with *P*(*i*|*i* < *k*_*c*_) the probability to be in state *i*, given that *i* < *k*_*c*_. Bayes’ law can be applied to find *P*(*i*|*i* < *k*_*c*_):

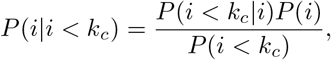

with:

- *P*(*i*) = *π*(*N* − *i* + 1), where we reversed the order in the steady-state distribution (2). Indeed, note that *i* is the initial state at the moment of the attention switch, i.e., when being in the steady-state regime from right before the switch, where state 1 was the target state (the reverse of Fig. 2).
- 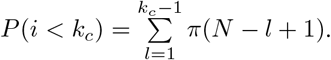
- *P*(*i* < *k*_*c*_|*i*) = 1 when *i* < *k*_*c*_ and = 0 otherwise.

Plugging this into (9) and using the definition of the MHT in (7) and the steady-state distribution in (2), we eventually find:

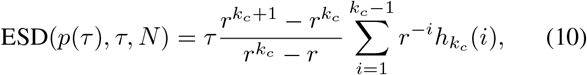

where 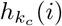 is given by (8). Note that ESD(*p*(*τ*), *τ*, *N*) (10) implicitly depends on *N* as the state index *k*_*c*_ = *c*(*N* − 1)+1 depends on *N*.

Given the *p*(*τ*)-performance curve of an AAD algorithm, constructed by piecewise linear interpolation through the points (*τ*_*i*_, *p*_*i*_), *i* ∈ {1, …, I} on the *p*(*τ*)-performance curve for which the AAD performance is evaluated on real data^5^, the optimal working point (*τ*_*opt*_, *p*_*opt*_) is defined as the pair for which the ESD(*p*(*τ*), *τ*, *N*) is minimal, given that *N* obeys the constraints of Section II-C3.

### E. The minimal expected switch duration

Optimizing *N*, *τ* and *p* now results in an optimal Markov chain that satisfies the stability constraints and has minimal ESD. The minimal ESD over the *p*(*τ*)-performance curve, which gave rise to the optimal working point (*τ*_*opt*_, *p*_*opt*_), can now be used as a single-number metric, referred to as the *minimal expected switch duration* (MESD), allowing to compare different AAD algorithms or parameter settings of the latter. This metric is defined as follows:

##### Definition (Minimal expected switch duration)

The minimal expected switch duration (MESD) is the expected time required to reach a predefined stable working region defined via the comfort level *c*, after an attention switch of the hearing aid user, in an optimized Markov chain as a model for an adaptive gain control system. Formally, it is the expected time to reach the comfort level *c* in the fastest Markov chain with at least *N*_min_ states for which 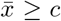, i.e., the lower bound 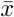 of the *P*_0_- interval is above *c*:

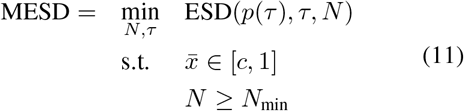

where ESD(*p*(*τ*), *τ*, *N*) is defined in (10) and 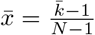 with 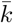 defined in (4).

The solution of optimization problem (11) is straightforward, given that ESD(*p*(*τ*), *τ*, *N*) is monotonically nondecreasing with *N* (see proof in Appendix E) for a fixed *τ*. Therefore, for each *τ*, choose the minimum 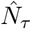 such that the two inequality constraints of (11) are obeyed (in Appendix C, it is proven that such an 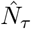 can always be found). As such, *N* is removed from the optimization problem, resulting in an unconstrained optimization problem:

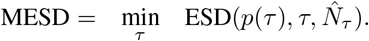

The MESD is then defined as the minimal ESD over all window lengths *τ*, at optimal working point (*τ*_*opt*_, *p*_*opt*_). Algorithm 1 summarizes the computation of the MESD metric.

Note that it is important to minimize the ESD over an as large as possible interval of decision window lengths, especially towards small decision window lengths. As a rule of thumb, assuming a monotonically decreasing accuracy with decreasing decision window length, one should consider a larger evaluation interval when the optimal working point (*τ*_*opt*_, *p*_*opt*_) obtained in Algorithm 1 is located at the boundary of the evaluated interval.

As an inherent by-product of the optimization problem in (11), the MESD metric also results in an optimal adaptive gain control system - optimal number of gains *N* and optimal working point (*τ*_*opt*_, *p*_*opt*_) - for a neuro-steered hearing aid.

## III. Experiments

We illustrate the MESD by applying the AAD algorithm in [6] on a dataset used in previous studies [10], which consists of 72 minutes of recorded EEG and audio data per subject (16 normal hearing subjects), who were instructed to listen to a specific speech stimulus in a competing two-speaker situation, including 24 minutes of repetitions but without intertrial attention switches. More details can be found in [10]. The 64-channel EEG data are bandpass filtered between 1 Hz and 9 Hz and downsampled to 20 Hz. The speech envelopes are computed using a power-law operation with exponent 0.6 after subband filtering [10] and are afterwards similarly bandpass filtered and downsampled. We assume that the clean envelopes of the original speech signals are available. In a practical hearing aid setting, these envelopes need to be extracted from the microphone recordings [12], [15], [16].

A linear spatio-temporal decoder, where the temporal dimension of the filter covers from 0 to 250 ms post-stimulus, is trained to decode the attended speech envelope from the EEG data by minimizing the mean-squared-error (MMSE) between the actually attended and predicted speech envelope on a training set. Per-subject decoders are trained and tested in a leave-one-trial-out fashion, using trials of consistent attention with a length of 60 s. Note that we apply the same adaptations to [6] as in [10], by training one decoder over all training trials and not averaging per-trial decoders. At test time, the trained filter decodes a speech envelope from a decision window of left-out EEG data of length *τ* (which is a subset of the leftout 60 s trial). The Pearson correlation coefficient is computed between the predicted speech envelope and the envelopes of both signals presented to the subject. The speech stream with the highest correlation is identified as the attended speaker.

To evaluate the algorithm on smaller decision window lengths, the left-out trial is segmented into smaller decision windows on which the corresponding decoder is applied. Reusing the decoders allows for fair comparison of the algorithm over different decision window lengths. The percentage of correct decisions *p*, per subject and decision window length *τ*, is computed as the total number of correct decisions divided by the total number of decisions over all trials.

### A. Hyperparameter choice

The MESD depends on three hyperparameters: the confidence level *P*_0_, the lower bound of the desired operating region *c* and the minimum number of states *N*_min_. When optimizing the design of a gain control system, the values of these hyperparameters can be set in a user-dependent fashion according to the needs and hearing capabilities of individual users (in particular for the desired comfort level *c*, which is very personal). However, in order to use the MESD as a standardized performance metric for comparing AAD algorithms, we also determined reasonable values for these hyperparameters and propose them as fixed inherent parameters of the MESD performance metric as a standard for future AAD algorithm comparison. We already motivated the choice for *N*_min_ = 5 in Section II-C2.

In order to find a value for the comfort level *c*, we need to determine the SNR (between attended and unattended speaker) corresponding to relative gain level *x* = 1 (SNR_max_) and the SNR corresponding to relative gain level *x* = *c*(SNR_*c*_). Using that *x* = 0.5 corresponds to 0 dB, *x* = *c* can be found from:

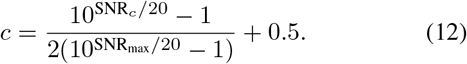

We here define SNR_max_ objectively as the speech reception threshold (SRT), corresponding to the 50% speech intelligibility level of the suppressed speaker, which should enable the hearing aid user to understand the suppressed speaker sufficiently, in order to assess whether (s)he wants to switch attention. Correspondingly, we define SNR_*c*_ as the SNR where there is full speech understanding *and* where the listening effort saturates, i.e., a higher SNR does not result in a better speech understanding nor less listening effort. In [23], the authors investigated the correct sentence recognition scores and peak pupil dilation, which quantifies the listening effort, when listening to standard Dutch sentences in the presence of a competing talker masker at SNR’s corresponding to daily life conditions. For normal hearing subjects, in their test setup, the average SRT corresponded to −11.2 dB (see Table 1 in [23]), such that SNR_max_ = 11.2 dB (as SNR_max_ is defined from the perspective of the attended, dominantly amplified speaker), while the correct sentence recognition score and listening effort saturate around 5 dB (see Fig. 1 in [23]). Plugging both values into (12) results in *c* = 0.65. We performed an additional subjective listening test on a story stimulus, which confirms that this is also a representative value for connected discourse stimuli (details on this experiment can be found in the supplementary material).

**Algorithm 1.**
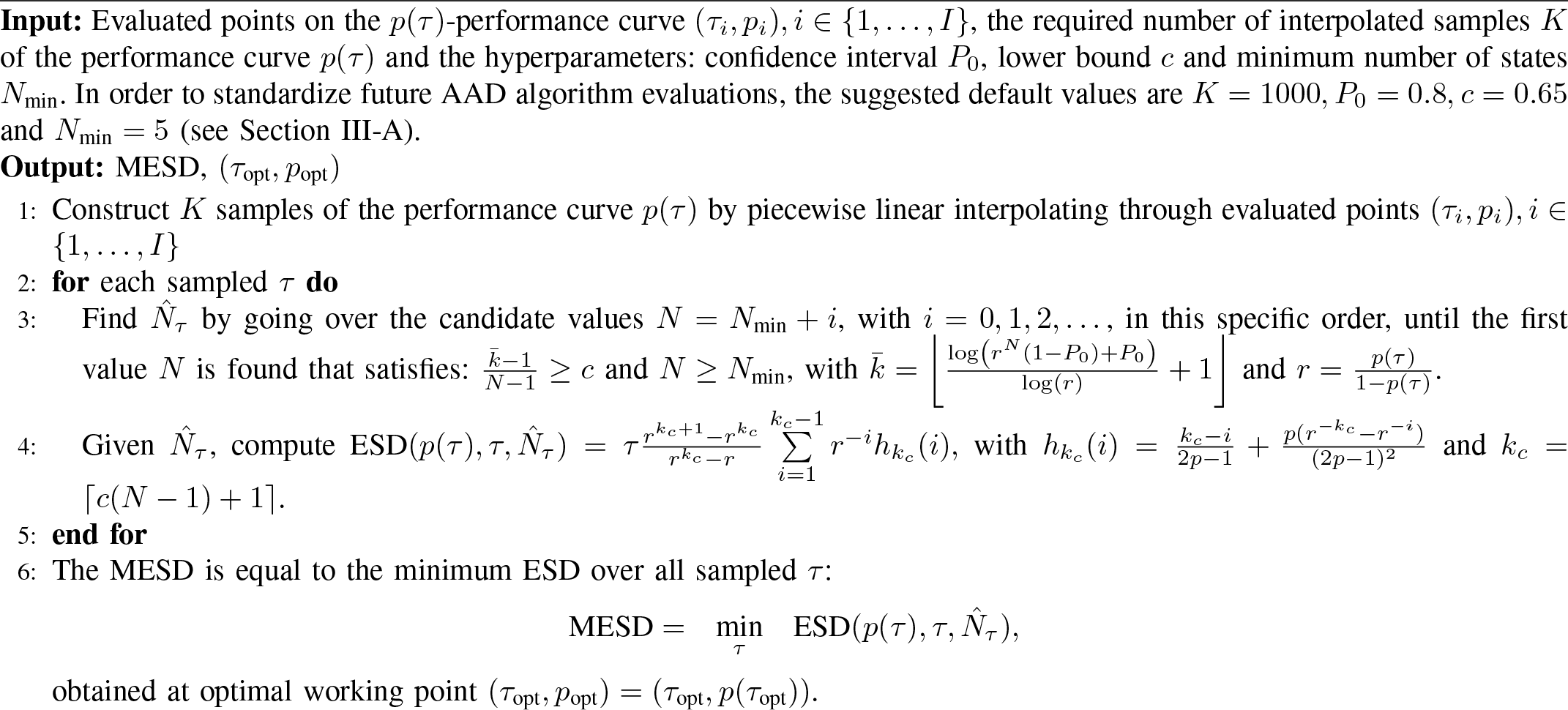
Computation of the MESD metric (code available in MESD toolbox)

Correspondingly, we choose *P*_0_ = 0.8, i.e., we require the system to be in the ‘comfortable’ operating region for 80% of the time. This confidence level yields a good trade-off between a high confidence level and a small enough MESD. Larger confidence levels result in a steep increase in MESD, yielding very high switch durations that are impractical, due to an overly strict confidence requirement.

A graphical analysis of the influence of the hyperparameters on the MESD metric is given in the supplementary material.

### B. Illustrative example: MESD-based performance evaluation

To illustrate why and how the MESD is useful in the evaluation of AAD algorithms, we apply it to an illustrative example in which we compare two variants of the MMSE decoder for AAD as proposed in [6] and [10], respectively.

#### 1) Description of the two variants

Given a training set of *M* data windows, in the first variant of [6] (also adopted in, e.g., [8]), per-window (corresponding to decision window length *τ*) decoders are computed, after which the *M* decoders are averaged to obtain one final decoder. The second variant of [10] (also adopted in, e.g., [12], [17], [18]), first averages the *M* per-window autocorrelation matrices (or equivalently: the windows are all concatenated) to train a single decoder across all training windows simultaneously. Similarly to [10], L_2_-norm regularization is added to the former method, in order to avoid overfitting effects due to the small amount of data per decoder. Note that no regularization is needed in the latter method because more data are used to train the decoder [10]. The decoders are again cross-validated in a leave-one-trial-out manner and the decoding accuracy is registered per regularization constant (between 10^−5^ and 10^2^, relative to the mean eigenvalue of the EEG autocorrelation matrix), for every decision window length. Again, the leave-one-trial out is done based on 60 s-trials, in order to keep the amount of training data constant for all decision window lengths. These trials are segmented in smaller windows when the decision window length decreases. Finally, for every window length *τ*, the maximum decoding accuracy in function of the regularization parameter is kept. Note that both variants thus use overall the same large amount of training data for each decision window length. When using a smaller decision window length, the decoders do not change for averaging autocorrelation matrices (as all data can be concatenated and the cross-validation is always done based on 60 s-trials), while for averaging decoders, more decoders are averaged out, each trained with a smaller amount of data.

#### 2) Subject-averaged comparison

The accuracies are averaged over all 16 subjects, resulting in one performance curve per variant, shown in Fig. 4 (with the standard deviation indicated by the shading). These performance curves can be interpreted in two ways, leading to two different conclusions depending on where we look. When looking at the region where *τ* > 30 s, one could conclude that both methods perform equally. This is because enough data are still used in the estimation of the per-window decoders in the method of [6]. However, in the region where *τ* < 30 s, one could conclude that averaging autocorrelation matrices is superior to averaging decoders, although, in total, an equal amount of training data has been used. Here, the loss of information when estimating decoders on small windows is not appropriately compensated by the averaging of a large number of decoders. Based on this analysis, it is not clear what the proper conclusion is, as it is a priori not clear what decision window lengths are more relevant in an AAD-based adaptive gain control system.

Here, the MESD and the corresponding optimal working point can resolve the dilemma mentioned above. Averaging autocorrelation matrices leads to an optimal Markov chain of seven states (optimized as in Section II-C and II-D), achieved at optimal working point (*τ*_*opt*_, *p*_*opt*_) = (2.54 s, 0.62), where the ESD is minimal. Taking a lower accuracy and decision window length would result in more states (see Section II-C3), which is not compensated by the smaller decision window length, resulting in a larger ESD. The number of states could be further minimized to five by increasing the decision window length, but the small decrease in target state *k*_*c*_ from five to four does not compensate enough for the increase in the decision window length. More details can be found in the supplementary material. A different optimal working point is chosen by the MESD metric for the case of averaging decoders, namely (*τ*_*opt*_, *p*_*opt*_) = (11.28 s, 0.68), meaning that it chooses for a slower, but more accurate decision process. Nevertheless, the MESD focuses in both cases on the smaller decision window lengths, based on a relevant and realistic criterion and thus overcomes potential inconclusiveness. It points at averaging autocorrelation matrices as a better way of computing the MMSE decoder, as it allows users to switch almost three times as fast.

#### 3) Statistical comparison

Instead of analyzing a single performance curve by averaging the performance curves per subject, which has the advantage of resulting in a *single*, generally optimal Markov chain and an easy-to-interpret over-all picture of the performance, one could also first compute the MESD per subject and perform a comparison based on these MESD performances using proper statistical testing procedures. A key aspect is that the MESD is a single-number metric, thereby allowing to straightforwardly perform statistical tests, while inherently taking the accuracy vs. decision window length trade-off into account. A paired, one-sided non-parametric Wilcoxon signed-rank test shows that the averaging of decoders significantly performs worse than the averaging of autocorrelation matrices (*W* = 0, *n* = 16, *p*-value < 0.001). This confirms the conclusion of [10], but more firmly, as we focused on the impact on a gain control system instead of arbitrarily choosing a decision window length to evaluate the related accuracy.

### C. Comparison of ITR_W/N_ and MESD

Similarly to the ITR_W/N_, the MESD quantifies the combination of the accuracy and decision time (window length) of an algorithm. As advocated before, the MESD uses, by design, a more relevant criterion to optimize the decision window length and accuracy in the context of AAD algorithms for gain control in hearing aids. By taking the maximum ITR_W/N_ (max-ITR) over all decision window lengths, one can define an alternative single-number metric (albeit less interpretable than the MESD). There is, however, a clear quantitative nonlinear relation between both metrics (Fig. 5a). Both the maximum of the Wolpaw ITR_W_ (1) (blue) and Nykopp ITR_N_^6^ (orange) are shown. Per subject, the performances are evaluated using MMSE decoders with averaging of autocorrelation matrices. Due to the nonlinearity, a significant difference in the max-ITR_W/N_ does not automatically imply a significant difference in MESD (and vice versa).

**Fig. 5:**
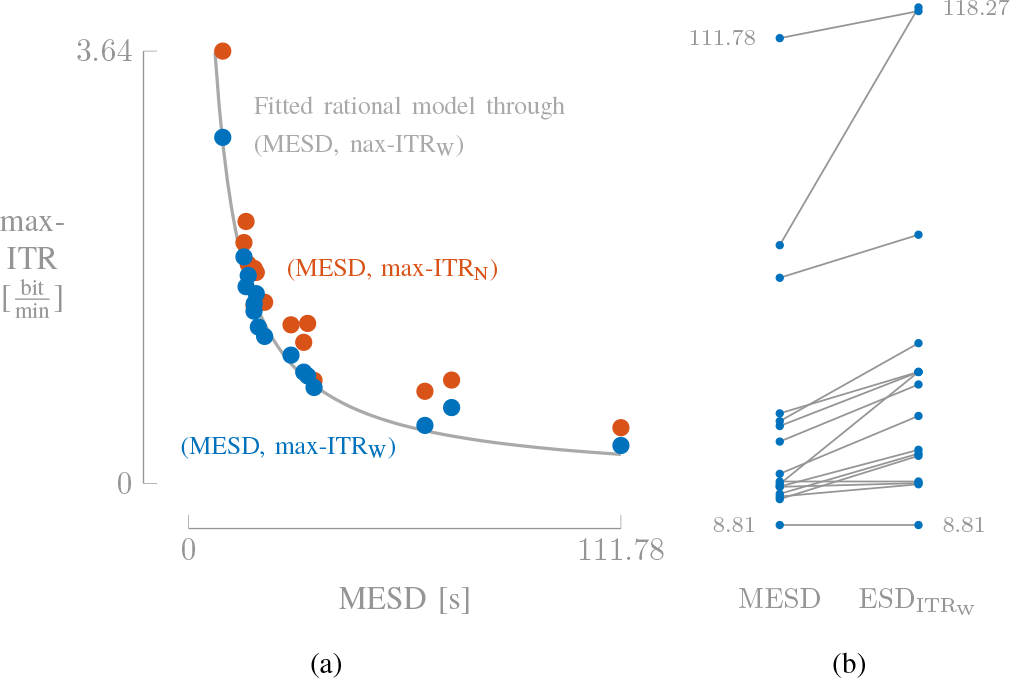
(a) A fitted rational model 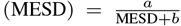 shows that there is a nonlinear relationship between the max-ITR_W_ and MESD. (b) Minizing the ESD (MESD) results in a significantly lower switch duration than optimizing the ESD based on the max-ITR_W_ (ineq), indicating that the MESD and ITR_W_ quantify performance in a fundamentally different way.

To highlight the differences between both metrics, we also compare the ESD, using the optimal working point based on maximizing the ITR_W_ 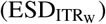, with the MESD (thus minimizing the ESD). Fig. 5b shows the per-subject differences in switch duration between the original MESD and the 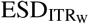 (a similar experiment can be conducted for ITR_N_). Note that for the majority of the subjects, there is a clear increase in switch duration, which already indicates that the ITR_W_ criterion does not select a working point on the *p*(*τ*)-performance curve that leads to an optimal working point for an adaptive gain control system, and therefore is not a representative metric to evaluate AAD algorithms in the context of, e.g., neuro-steered hearing aids. Moreover, several relative differences between subjects have changed, indicating that both criteria fundamentally differ. A non-parametric Wilcoxon signed-rank test (*W* = 0, *n* = 16, *p*-value < 0.001) confirms that there is significant difference between both switch durations. Optimality in case of ITR_W_ thus has a fundamentally different and less clear interpretation than in case of the MESD, which stems from the fact that ITR_W/N_ focuses on optimizing information transfer rate as such, which is different from optimizing and stabilizing a gain control system.

In conclusion, it is more relevant to perform (statistical) analysis on a metric that represents a major goal in the context of hearing aids: fast, accurate and *stable* switching.

## IV. Conclusion

In this paper, we have developed a new *interpretable* performance metric to evaluate AAD algorithms for AAD-based gain control: the minimal expected switch duration. This metric quantifies the expected time to perform a gain switch after an attention switch of the user in an AAD-based adaptive gain control system, towards a comfort level (*c* = 0.65) that can be maintained for at least 80% of the time. It is based on the concept of the mean hitting time in a Markov chain model, which resulted in a closed-form expression because of the specific line-graph structure. The MESD can be computed from the performance curve of an AAD system by minimizing the expected switch duration over this curve, after designing an optimal Markov chain such that it is for *P*_0_ = 80% of the time in an optimal operating region. The derivation of the MESD also results in a design methodology for an optimal AAD-based volume control system, as a by-product. The fact that the MESD provides a single-number AAD performance metric, that combines accuracy and decision window length and that is also interpretable and relevant within the context of neuro-steered hearing prostheses, is paramount in order to uniformize the evaluation of AAD algorithms in this context.

Experiments on real EEG and audio data showed that this metric can be used to globally compare AAD systems, both between subjects and between algorithms. Finally, we showed that the MESD is quantitatively related to the ITR_W/N_, but that it uses a fundamentally different criterion that is more relevant in the context of hearing aids.

As a final remark, note that this metric can be easily extended to other BCI applications. In, for example, 1D cursor control using EEG (e.g., [25]), it could be used to quantify the expected time needed to move a cursor or object from one end to the other end in a stable fashion.

## Appendix A

The steady-state distribution can be found from the global balance equations and the normalization condition [26]:

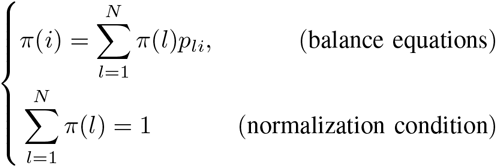

where *p*_*li*_ corresponds to the transition probability from state *l* to state *i*. We can solve the balance equations recursively, starting from *π*(1):

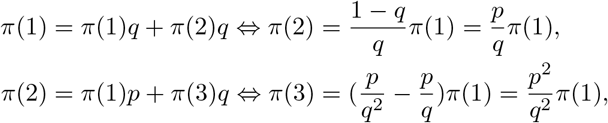

By working out the recursion further on and by defining 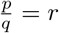, it can be seen that:

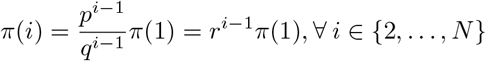

*π*(1) can be found from the normalization condition:

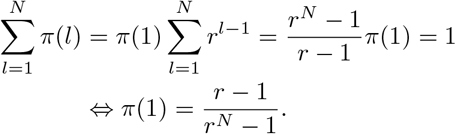

## Appendix B

Starting from (3) and using the steady-state distribution in (2), we obtain:

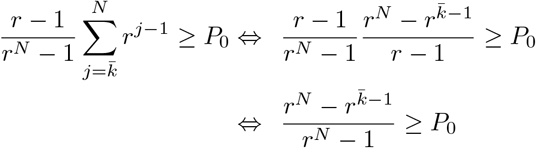

Since we assume that *p* > 0.5, it holds that *r* > 1. Hence, both the numerator and denominator are positive. Furthermore, the log-function is a monotonically increasing function, such that it can be applied to both sides without changing the inequality:

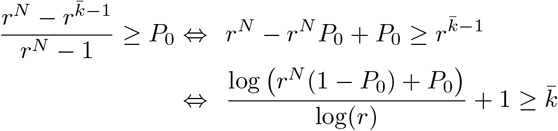

Flooring the last expression leads to (4).

## Appendix C

In this appendix, we prove that there always exists a solution for *N* such that (6) is satisfied. Using (4), it can be seen that:

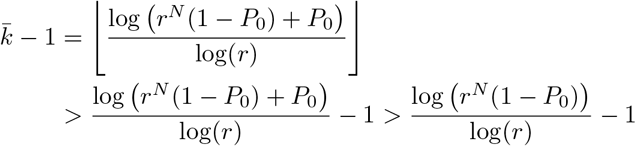

such that the constraint (6) is always satisfied when

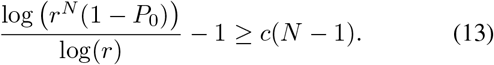

Solving for *N* yields:

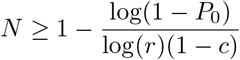

## Appendix D

The MHT can be found from the recursive definition in [26]:

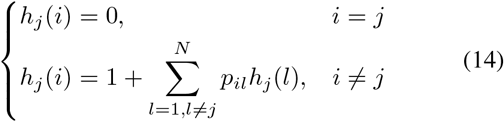

When *i* ≤ *j*, *h*_*j*_(*i*) can be found by starting the recursion in (14) with *h*_*j*_(1):

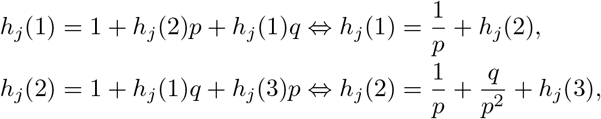

Eventually, it can be found that:

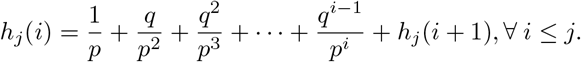

For *i* = *j −* 1, this results in:

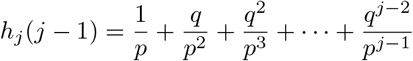

where *h*_*j*_(*j*) = 0 because of (14). By propagating the solutions backward, we find:

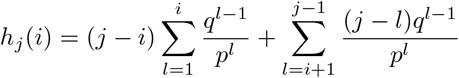

By computing the sums and simplifying the expressions, the expression in (8) is found.

## Appendix E

We prove that ESD(*p*, *τ*, *N*) in (10) is monotonically non-decreasing with *N*. Starting from (9) and using Bayes law as in the manuscript, the ESD can be written as:

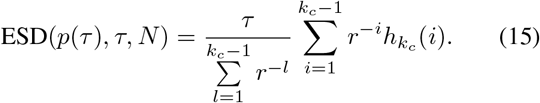

ESD(*p*(*τ*), *τ*, *N*) only implicitly depends on *N* via *k*_*c*_ = ⌈*c*(*N* − 1) + 1⌉. We use the notation *k*_*c*_(*N*) to explicitly show that *k*_*c*_ is a function of *N*. Note that *k*_*c*_(*N* + 1) ≤ *k*_*c*_(*N*)+ 1 as *k*_*c*_(*N* + 1) = ⌈*cN*⌉ + 1, while *k*_*c*_(*N*)+ 1 = ⌈*cN* + 1 *c* + 1⌉ ≥ ⌈*cN*⌉ + 1 as *c* ≤ 1. Furthermore, *k*_*c*_(*N*) is monotonically increasing with *N*. This means that there are two possibilities: when *N → N* + 1, then either *k_c_ → k*_*c*_ or *k*_*c*_ → *k*_*c*_ + 1.

- *Case k_c_ → k*_*c*_: from (15) it can be easily seen that in this case ESD(*p*(*τ*), *τ*, *N* + 1) = ESD(*p*(*τ*), *τ*, *N*), as 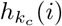 (8) only depends on *k*_*c*_ and not explicitly on *N*.
- *Case k*_*c*_ → *k*_*c*_ + 1: the proof boils down to proving that:

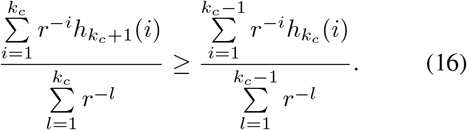

If we can show that ∀ *i* ≤ *k*_*c*_ − 1:

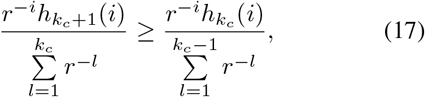

then (16) is true (note that 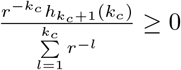. From (8) it can be found that:

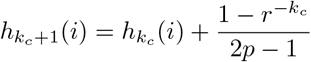

By using the previous result and substituting 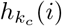 with (8) in (17), we eventually find after some straight forward algebraic manipulations that (17) boils down to:

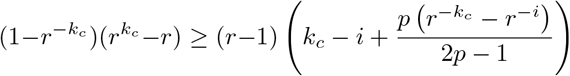

After some further manipulation and using 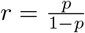, this becomes:

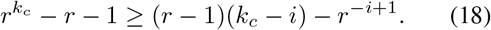

We now show that the right-hand side of (18) is a decreasing function with *i ≤ k_c_ −* 1. If *f*(*i*) = (*r −*1)(*k_c_ − i*) *− r^−i^*^+1^, then *f*(*i* + 1) is equal to:

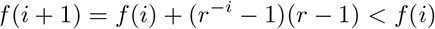

because *r* > 1 and *i* ≥ 1. Given that the right-hand side of (18) is decreasing with *i*, we only have to proof (18) for *i* = 1:

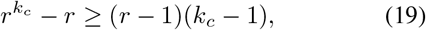

which can be easily proven by induction. For *k*_*c*_ = 2 it holds that:

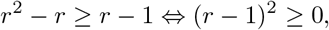

which is evidently true. Now we prove that, if (19) is true for *k*_*c*_ = *j* ≥ 2, then it is also true for *k*_*c*_ = *j* + 1. Setting *k*_*c*_ = *j*, (19) can be rewritten as

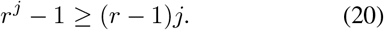

Furthermore, since *r* > 1, we have that *r^j^*^+1^ − *r* ≥ *r*^*j*^ − 1 and therefore (19) holds for *k*_*c*_ = *j* + 1, using the induction hypothesis in (20). This concludes the proof.

## Supplementary Material

In the supplementary material, related to the paper *An Interpretable Performance Metric for Auditory Attention Decoding Algorithms in a Context of Neuro-Steered Gain Control*, we describe a subjective listening test to validate the choice for the comfort level *c* = 0.65 (Section I) and elaborate on the influence of the hyperparameters *P*_0_ (the confidence level) and *c* (comfort level) on the MESD metric (Section II). Furthermore, we investigate in Section III how the ESD and the number of states of the optimized Markov chain depend on the decision window length and accuracy, for the MMSE-based decoder with averaging of autocorrelation matrices.

### I. Validation of the comfort level *c*

To validate the chosen *c*-value (*c* = 0.65) of Section III-A in case of a (more relevant) connected discourse stimulus instead of standard sentences (as used in Section III-A), we conducted a subjective listening experiment to determine SNR*_*c*_*. Eight normal hearing participants, aged between 24 and 29 and with Dutch as their mother tongue, were asked to listen to a mixture of two non-standardized, commercial recordings of stories, 6 min and 34 s long. The stimuli were biologically calibrated. The participants were allowed to adapt the SNR with a slider between 0 and 50 dB and were instructed to select the minimal SNR (between the dominantly amplified speaker and the competing speaker) that still allowed them to comfortably listen to the dominantly amplified speaker for a duration of, e.g., 30 min. When they selected a value for SNR*_*c*_*, they were instructed to listen to the dominantly amplified speaker for three more minutes at their selected SNR*_*c*_*, where now the previously suppressed speaker is the dominantly amplified speaker. As a validation procedure, the participants self-reported their listening effort, probing the amount of effort required to understand the loudest speaker. A review on the self-reported listening effort and other methods to assess listening effort can be found in [1]. The minimal reported, maximal reported and median SNR_*c*_ is equal to 4.56 dB, 23.55 dB and 10.89 dB. All reported listening efforts were below 25%.

To obtain the SRT, we used the results from [2], where they performed a similar experiment (using similar conditions) in an age-matched, normal hearing group to determine the SRT of connected discourse using the self-assessed Békesy procedure. We use the median SRT = −16.27 dB as a value for SNR_max_ = 16.27 dB. Note that this SRT differs from the one reported in Section III-A, as we are now dealing with a connected discourse instead of standard sentences, while also a different procedure for assessing speech intelligibility has been used.

The resulting *c*-value (12) is equal to *c* = 0.727. Given the large variability on the reported comfort level, we consider this value to be reasonably close to the proposed value *c* = 0.65, which was calculated based on data from the literature.

### II. The relation between the MESD and the hyperparameters

Fig. 1 shows how the MESD metric depends on the hyperparameters *P*_0_ (the confidence level) and *c* (the comfort level). The MESD’s are based on the results of an MMSE-based decoder with averaging of autocorrelation matrices, described in Section III and Fig. 4 in the original paper. When varying one hyperparameter, the other hyperparameters are kept constant at their default values (*P*_0_ = 0.8*, c* = 0.65*, N*_min_ = 5). The black diamonds indicate the chosen hyperparameter value in the paper. Fig. 1a shows that *P*_0_ = 0.8 yields a good trade-off between a high confidence level and a small enough MESD. As the MESD has a positive second-order derivative in function of *P*_0_, an extra amount of confidence results in an even larger increase in MESD, which is why it is important to choose its value as low as possible, without giving too much in on the reliability of the gain control system.

The MESD is a discrete function of the comfort level *c* (Fig. 1b). As the lower bound of the *P*_0_-confidence interval needs to be above comfort level *c*, a higher comfort level results in more states and thus in a higher MESD. Note that because of the flooring operation in (4), this a discrete function. Again, higher comfort levels result in a steeper increase in switch duration. The comfort level *c* = 0.65 that resulted from the analysis and experiments in Section III-A of the paper and Section I of the supplementary material seems to avoid this high cost of extra comfort while assuring, by design, enough comfort for the user.

### III. THE ESD and number of states in function of the decision window length

In Section III-B, the MESD has been applied to the performance curve of the MMSE-based decoder with averaging of autocorrelation matrices versus averaging decoders (Fig. 4 in the original paper). We mentioned that the optimal MESD for averaging of to correlation matrices is obtained at a Markov chain of seven states. Fig. 2 shows the optimal number of states 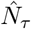 and target state *k*_*c*_ per decision window length (see Section II-E and Algorithm 1) and the ESD per decision window length, at the optimal number of states 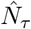. It is over this curve that the ESD is minimized to obtain the MESD (Section II-E and Algorithm 1).

**Fig. 1:**
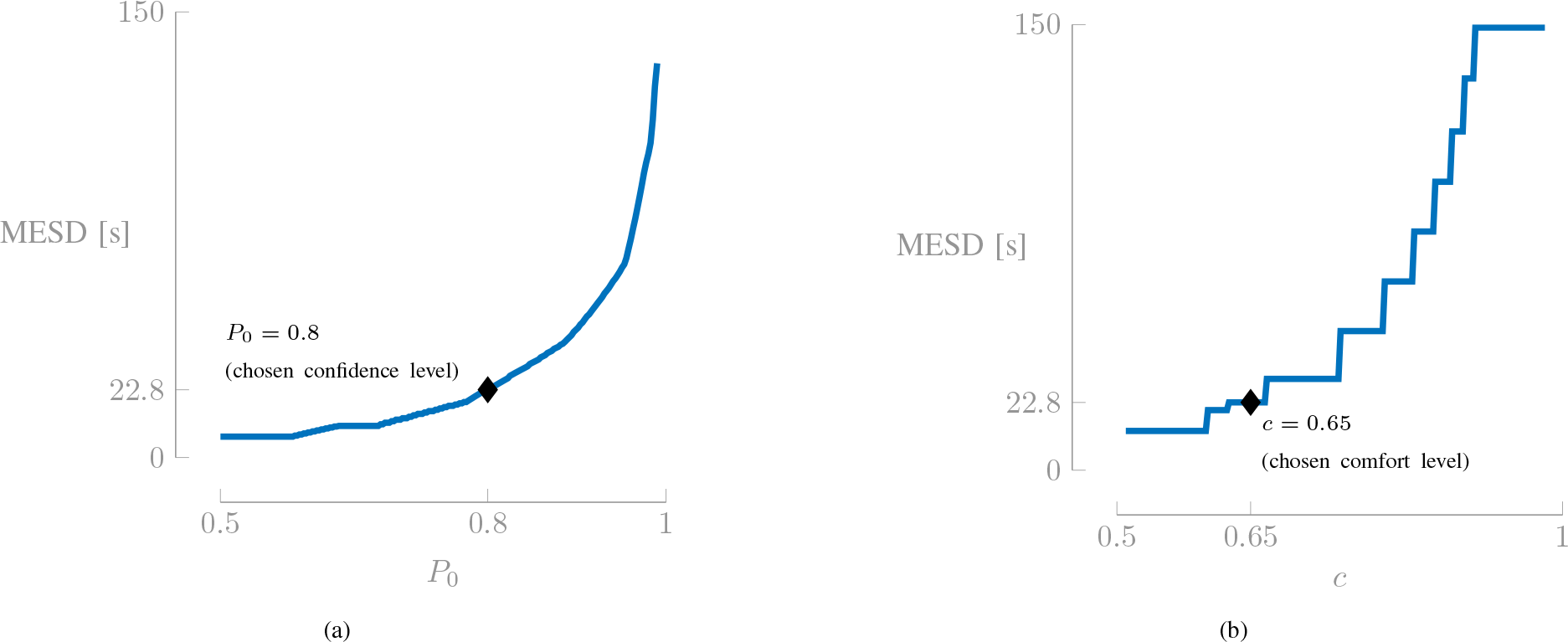
The MESD increases in function of (a) the confidence level *P*_0_, with a positive second-order derivative and (b) the comfort level *c*, in a discrete way, also with an increasing slope. The MESD’s are shown for the performance curve of the MMSE-based decoder with averaging of autocorrelation matrices. The chosen confidence level and comfort level are indicated by a diamond (♦). When varying a hyperparameter, the other hyperparameter is kept constant at the default value (*c* = 0.65*, P*_0_ = 0.8).

In Fig. 2, it can be seen that when 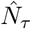 remains constant, the ESD increases almost linear with decision window length *τ*. In (10), when the number of states *N* and thus target state *k*_*c*_, remains constant, it appears that the step time *τ* is the dominant factor over the variation in transition probability *p*. This implies that the interesting decision window lengths coincide with changes in the number of states. Relative to 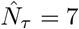 at the MESD, an increase in decision window length results in a decrease of 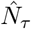 to five. However, the target state *k*_*c*_ only decreases from five to four, such that the drop in ESD around ≈ 6 s is not large enough to decrease below the minimal ESD for seven states. When decreasing *τ*, 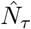 and *k*_*c*_ increase steeply because of the steep decrease in accuracy (Fig. 4 in the original paper), which is not sufficiently compensated by the small decrease in step time *τ*. The AAD accuracy *p* (depending on decision window length *τ*) thus mainly plays a role in determining the optimal number of states 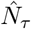 via the design constraints (Section II-C), which is the first step in optimizing the ESD (Section II-E and Algorithm 1), while the transition points of 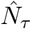 are most interesting for minimizing the ESD to obtain the MESD, as the ESD almost linearly increases with *τ* for a constant 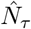.

**Fig. 2:**
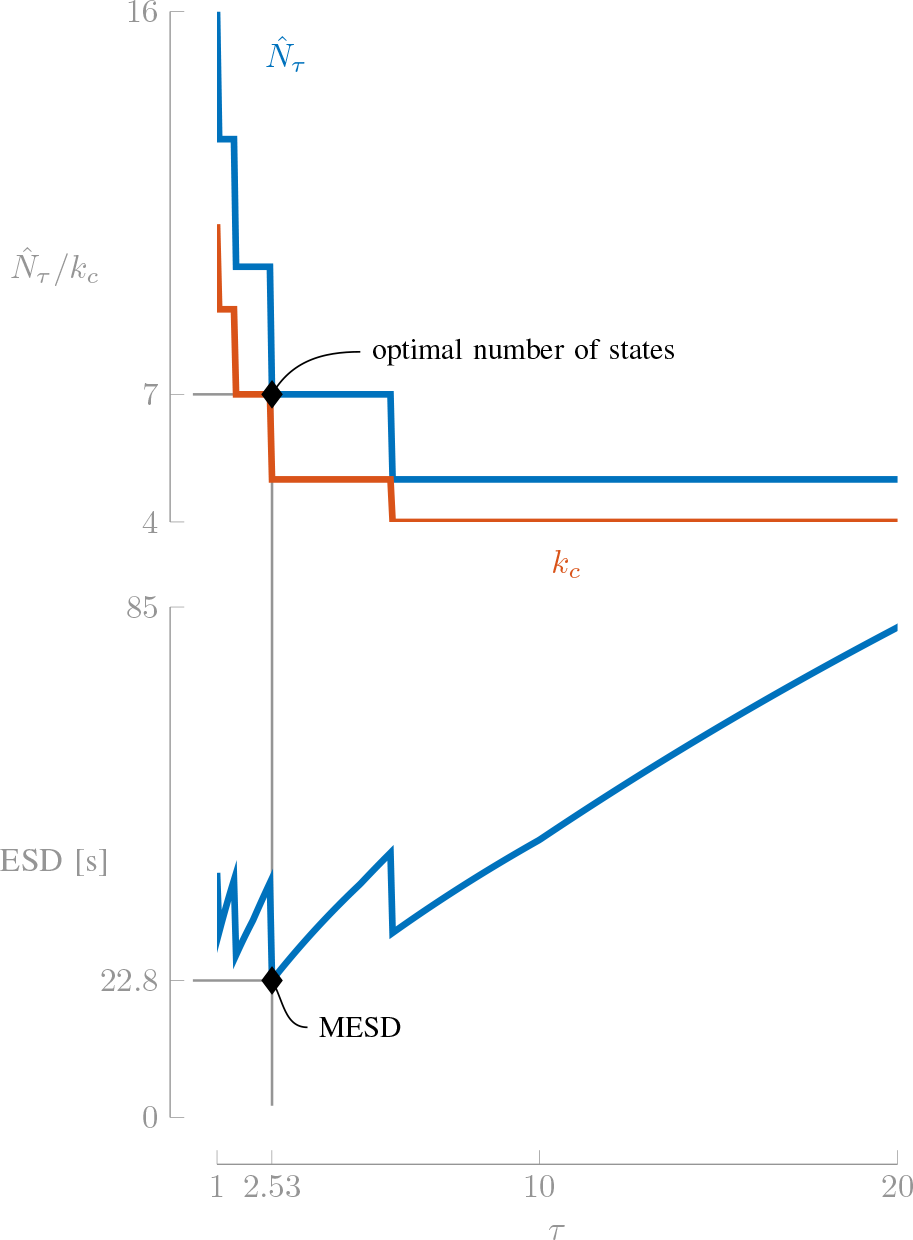
The optimal number of states 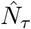 and corresponding target state *kc* decrease in function of the decision window length *τ*. The minimal ESD (MESD) depends both on the optimal number of states (via the AAD accuracy) and the decision window length.

1 We provide an open-source implementation to compute the MESD metric, which can be found online on https://github.com/exporl/mesd-toolbox.

2 It is noted that the ITR metric uses a similar assumption, as it implicitly assumes independence between consecutive messages (i.e. AAD decisions).

3 Note that due to the discretization of *x*, the probability of being in 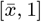 is generally larger than *P*_0_. However, (4) ensures that 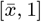 is the *smallest* possible interval such that 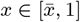 for at least *P*_0_ percent of the time.

4 This holds unless *p* > *P*_0_, in which case the *P*_0_-confidence interval collapses to a single state *N* = 4.

5 In this paper, we assume that *p* is fixed and evaluated over all data windows (batch)

6 The COCOHA MATLAB toolbox [24] has been used to compute ITR_N_.

## References

[1] E. C. Cherry, “Some Experiments on the Recognition of Speech, with One and with Two Ears,” J. Acoust. Soc. Am., vol. 25, no. 5, pp. 975–979, 1953.

[2] N. Mesgarani and E. F. Chang, “Selective cortical representation of attended speaker in multi-talker speech perception,” Nature, vol. 485, no. 7397, pp. 233–236, 2012.

[3] S. J. Aiken and T. W. Picton, “Human Cortical Responses to the Speech Envelope,” Ear and Hearing, vol. 29, no. 2, pp. 139–157, 2008.

[4] J. R. Kerlin, A. J. Shahin, and L. M. Miller, “Attentional Gain Control of Ongoing Cortical Speech Representations in a “Cocktail Party”,” J. Neurosci., vol. 30, no. 2, pp. 620–628, 2010.

[5] E. M. Z. Golumbic et al., “Mechanisms Underlying Selective Neuronal Tracking of Attended Speech at a “Cocktail Party”,” Neuron, vol. 77, no. 5, pp. 980–991, 2013.

[6] J. A. O’Sullivan et al., “Attentional Selection in a Cocktail Party Environment Can Be Decoded from Single-Trial EEG,” Cereb. Cortex, vol. 25, no. 7, pp. 1697–1706, 2014.

[7] T. de Taillez, B. Kollmeier, and B. T. Meyer, “Machine learning for decoding listeners’ attention from electroencephalography evoked by continuous speech,” Eur. J. Neurosci., 2017.

[8] E. Alickovic, T. Lunner, F. Gustafsson, and L. Ljung, “A Tutorial on Auditory Attention Identification Methods,” Front. Neurosci., vol. 13, p. 153, 2019.

[9] A. de Cheveigné et al., “Decoding the auditory brain with canonical component analysis,” NeuroImage, vol. 172, pp. 206–216, 2018.

[10] W. Biesmans, N. Das, T. Francart, and A. Bertrand, “Auditory-inspired speech envelope extraction methods for improved EEG-based auditory attention detection in a cocktail party scenario,” IEEE Trans. Neural Syst. Rehabil. Eng., vol. 25, no. 5, pp. 402–412, 2017.

[11] S. Miran et al., “Real-Time Tracking of Selective Auditory Attention from M/EEG: A Bayesian Filtering Approach,” Front. Neurosci., vol. 12, p. 262, 2018.

[12] S. Van Eyndhoven, T. Francart, and A. Bertrand, “EEG-Informed Attended Speaker Extraction From Recorded Speech Mixtures With Application in Neuro-Steered Hearing Prostheses,” IEEE Trans. Biomed. Eng., vol. 64, no. 5, pp. 1045–1056, 2017.

[13] J. O’Sullivan et al., “Neural decoding of attentional selection in multi-speaker environments without access to clean sources,” J. Neural Eng., vol. 14, no. 5, p. 056001, 2017.

[14] N. Das, S. Van Eyndhoven, T. Francart, and A. Bertrand, “EEG-based Attention-Driven Speech Enhancement For Noisy Speech Mixtures Using N-fold Multi-Channel Wiener Filters,” in Proc. Eur. Signal Process. Conf. (EUSIPCO). IEEE, 2017, pp. 1660–1664.

[15] C. Han et al., “Speaker-independent auditory attention decoding without access to clean speech sources,” Sci. Adv., vol. 5, no. 5, pp. 1–12, 2019.

[16] A. Aroudi and S. Doclo, “Cognitive-driven binaural LCMV beamformer using EEG-based Auditory Attention Decoding,” in Proc. IEEE International Conference on Acoustics, Speech and Signal Processing (ICASSP), 2019, pp. 406–410.

[17] N. Das, A. Bertrand, and T. Francart, “EEG-based auditory attention detection: boundary conditions for background noise and speaker positions,” J. Neural Eng., vol. 15, no. 6, 2018, 066017.

[18] A. Aroudi, B. Mirkovic, M. De Vos, and S. Doclo, “Impact of Different Acoustic Components on EEG-Based Auditory Attention Decoding in Noisy and Reverberant Conditions,” IEEE Trans. Neural Syst. Rehabil. Eng., vol. 27, no. 4, pp. 652–663, 2019.

[19] S. A. Fuglsang, T. Dau, and J. Hjortkjær, “Noise-robust cortical tracking of attended speech in real-world acoustic scenes,” NeuroImage, vol. 156, pp. 435–444, 2017.

[20] J. R. Wolpaw, H. Ramoser, D. J. McFarland, and G. Pfurtscheller, “EEG-Based Communication: Improved Accuracy by Response Verification,” IEEE Trans. Rehabil. Eng., vol. 6, no. 3, pp. 326–333, 1998.

[21] D. D. E. Wong et al., “A Comparison of Regularization Methods in Forward and Backward Models for Auditory Attention Decoding,” Front. Neurosci., vol. 12, p. 531, 2018.

[22] S. Geirnaert, T. Francart, and A. Bertrand, “A New Metric to Evaluate Auditory Attention Detection Performance Based on a Markov Chain,” in Proc. Eur. Signal Process. Conf. (EUSIPCO), September 2019, Accepted for publication.

[23] B. Ohlenforst et al., “Impact of stimulus-related factors and hearing impairment on listening effort as indicated by pupil dilation,” Hear. Res., vol. 351, pp. 68–79, 2017.

[24] D. D. E. Wong, J. Hjortkjær, E. Ceolini, and A. de Cheveigné, “COCOHA Matlab Toolbox,” https://cocoha.org/the-cocoha-matlab-toolbox/,v0.5.0, March 2018.

[25] G. Schalk et al., “BCI2000: A General-Purpose Brain-Computer Interface (BCI) System,” IEEE Trans. Biomed. Eng., vol. 51, no. 6, pp. 1034–1043, June 2004.

[26] P. Brémaud, Markov chains: Gibbs fields, Monte Carlo Simulation, and Queues, ser. Texts in Applied Mathematics. New York: Springer Science & Business Media, 2013, vol. 31.

## References

[1] R. McGarrigle, K. J. Munro, P. Dawes, A. J. Stewart, D. R. Moore, J. G. Barry, and S. Amitay, “Listening effort and fatigue: What exactly are we measuring? A British Society of Audiology Cognition in Hearing Special Interest Group ‘white paper’,” Int. J. Audiol., vol. 53, no. 7, pp. 433–445, 2014.

[2] L. Decruy, N. Das, E. Verschueren, and T. Francart, “The Self-Assessed Békesy Procedure: Validation of a Method to Measure Intelligibility of Connected Discourse,” Trends Hear., vol. 22, pp. 1–13, 2018.

